# Fiber dispersion in the right ventricle: A comparison of constitutive neural network predictions with experimental data

**DOI:** 10.64898/2026.05.11.724139

**Authors:** Parag S. Ingalkar, Sotirios Kakaletsis, Manuel K. Rausch, Ellen Kuhl, Denisa Martonová

**Affiliations:** Institute of Applied Mechanics, Egerlandstraße 5, Friedrich-Alexander-Universität Erlangen-Nürnberg, 91058 Erlangen, Germany; Department of Aerospace Engineering and Engineering Mechanics, The University of Texas at Austin, Texas 78712, USA; Department of Mechanical Engineering, 318 Campus Drive, Stanford University, California 94305, USA

**Keywords:** right ventricle, constitutive neural networks, model discovery, passive material model, fiber dispersion

## Abstract

The mechanical behavior of right ventricular (RV) myocardium is governed by its anisotropic microstructure, yet constitutive models that account for fiber dispersion and enable reliable parameter identification remain limited. In this study, we propose a physics-embedded constitutive neural network framework for automated discovery of strain energy functions and microstructural parameters from experimental data. The model is formulated within an incompressible, orthotropic hyperelastic setting using invariant-based representations. Fiber, sheet, and normal directions are incorporated through a rotated structural basis, and dispersion effects are modeled using a generalized structure tensor approach. The framework is trained on multi-axial mechanical data from ovine RV myocardium, including uniaxial tension–compression and simple shear tests. We investigate two training scenarios: (i) full datasets containing both tensile and compressive regimes and (ii) datasets restricted to tensile loading. In both cases, the model accurately reproduces the measured stress–strain responses and identifies sparse, interpretable constitutive models which involve isotropic, anisotropic, and coupling invariants. However, the identifiability of microstructural parameters strongly depends on the available loading conditions. While tensile-only data yield higher predictive accuracy, they result in non-unique or biased estimates of fiber dispersion. In contrast, inclusion of compressive data enables consistent identification of dispersion parameters by separating fiber and matrix contributions. These results highlight the importance of multi-axial loading data for robust parameter identification and demonstrate the capability of constitutive neural network-based approaches for data-driven modeling of anisotropic soft tissues.

## 1. Introduction

For decades, research in cardiac mechanics has focused primarily on the left ventricle (LV), while the right ventricle (RV) has received comparatively limited attention (Rigolin et al., 1995; Valdez-Jasso et al., 2012; Kakaletsis et al., 2021, 2023). Early physiological interpretations suggested that the RV played a secondary mechanical role in global circulation and thereby reinforced the perception that detailed structural and material characterization was less critical (Voelkel et al., 2006; Amsallem et al., 2018; Monteagudo-Vela et al., 2023). In addition, the RV’s thin wall, crescent-like geometry, and anterior anatomical position complicate both imaging and mechanical testing (Friedrich and Chetrit, 2023). Together, these challenges have delayed the development of a comprehensive mechanical description of RV myocardium.

This perspective has shifted substantially. Clinical evidence now demonstrates that RV performance is a key determinant of outcomes in pulmonary hypertension, right-sided myocardial infarction, biventricular heart failure, and respiratory disease. Importantly, the RV cannot be interpreted as a geometric variant of the LV. The two chambers differ markedly in wall thickness, loading environment, geometry, and microstructural organization. Studies of myocardial architecture have revealed distinct chamber-specific patterns of fiber orientation and transmural variation (Streeter et al., 1969; Holzapfel and Ogden, 2009), which emphasize the need for dedicated constitutive characterization of the RV. A rigorous understanding of its passive mechanical behavior is therefore essential for elucidating right ventricular function in both physiological and pathological settings.

Extensive investigations of LV mechanics have established a methodological foundation for myocardial constitutive modeling. Mechanical testing under planar and shear deformation modes (Dokos et al., 2002; Sommer et al., 2015), combined with structurally motivated anisotropic hyperelastic formulations (Holzapfel and Ogden, 2009; Guan et al., 2022; Martonová et al., 2021), has enabled systematic parameter identification and the comparison of competing constitutive laws. More recently, inverse finite element approaches have replaced simplified homogeneous assumptions in order to capture heterogeneous deformation fields observed in experiments and developments in imaging have facilitated incorporation of microstructural information into constitutive descriptions (Avazmohammadi et al., 2017; Kolawole et al., 2025; Martonová et al., 2025).

An important characteristic of myocardial tissue is its anisotropic architecture. Cardiomyocytes are organized into fibers with preferred orientations that vary transmurally and assemble into sheet-like laminae that contribute to complex deformation mechanisms. Constitutive models typically introduce local fiber, sheet, and sheet-normal directions to represent this orthotropic structure. However, many formulations assume perfectly aligned fiber families at each material point. This assumption neglects the inherent variability observed in biological tissues, where fibers are distributed around a mean orientation rather than being identically aligned. To address this variability, dispersion-based modeling frameworks have been proposed. Early approaches described fiber families as continuous distributions over orientation space (Lanir, 1983). While theoretically rigorous, full angular integration formulations are computationally demanding. Alternatively, generalized structure tensor models provide an efficient algebraic approximation of fiber dispersion effects and have been widely adopted for soft biological tissues (Gasser et al., 2006; Holzapfel et al., 2015). Although dispersion has been investigated in myocardium and related tissues, its quantitative influence on constitutive model identification, particularly in the right ventricle, remains incompletely characterized. Moreover, systematic shifts in mean fiber orientation relative to canonical directions may alter predicted mechanical responses in ways not captured by dispersion parameters alone.

These limitations motivate the use of data-driven constitutive approaches, which enable inference of material behavior directly from experimental observations without prescribing a fixed functional form. Physics-embedded neural networks enforce kinematic constraints, material symmetry, and thermodynamic consistency directly into their architecture and thereby enable automated discovery of admissible strain energy functions from experimental data (Linka et al., 2021, 2023; Kalina et al., 2021; Linden et al., 2023; Holthusen et al., 2024; Flaschel et al., 2025). Rather than prescribing a fixed analytical form, these models infer functional relationships directly from measurements while preserving physical consistency. Their application to biological tissues has demonstrated strong predictive capability in capturing complex anisotropic behavior (Linka et al., 2023; Martonová et al., 2024, 2025).

We address these open questions by integrating detailed experimental measurements from ovine right ventricular myocardium (Kakaletsis et al., 2021) with a physics-informed constitutive neural network framework. Specifically, we (i) analyze mechanical data obtained from ovine RV tissue, (ii) incorporate both fiber dispersion and systematic shifts in mean fiber angle into the constitutive description, (iii) train a constitutive neural network to autonomously discover both necessary material model terms and dispersion parameters directly from the mechanical response, (iv) compare the predicted dispersion parameters with independently measured structural data, and (v) evaluate whether this combined strategy yields an accurate and physically consistent material model for right ventricular myocardium.

This work links experimentally observed microstructure in ovine RV tissue with automated constitutive model discovery. It advances understanding of right ventricular mechanics and establishes a framework for integrating structural uncertainty into myocardial material modeling.

## 2. Continuum modeling approach

We model the right-ventricular myocardium as an incompressible, hyperelastic, orthotropic material (Kakaletsis et al., 2021). The constitutive response of the material is governed by the local deformation and the orientation of the underlying microstructural components (Holzapfel and Ogden, 2009). We denote by *φ* the deformation map from the reference configuration **X** to the current configuration **x** = *φ*(**X**). The deformation gradient and its Jacobian are

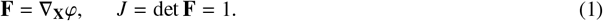

We multiply the deformation gradient by its transpose from the left to obtain the right Cauchy-Green deformation tensor **C** = **F**^T^**F**. We assume three orthogonal microstructural directions, the fiber, sheet, and normal directions ***f***_0_, ***s***_0_, ***n***_0_, defined in the reference configuration relative to the global basis {***e***_1_, ***e***_2_, ***e***_3_}. The fiber and sheet directions follow from an in-plane rotation in the *e*_1_*e*_2_ plane,

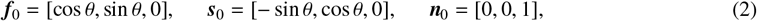

where *θ* denotes the mean fiber angle. The normal direction ***n***_0_ aligns with ***e***_3_, see Figure 1.

**Figure 1:**
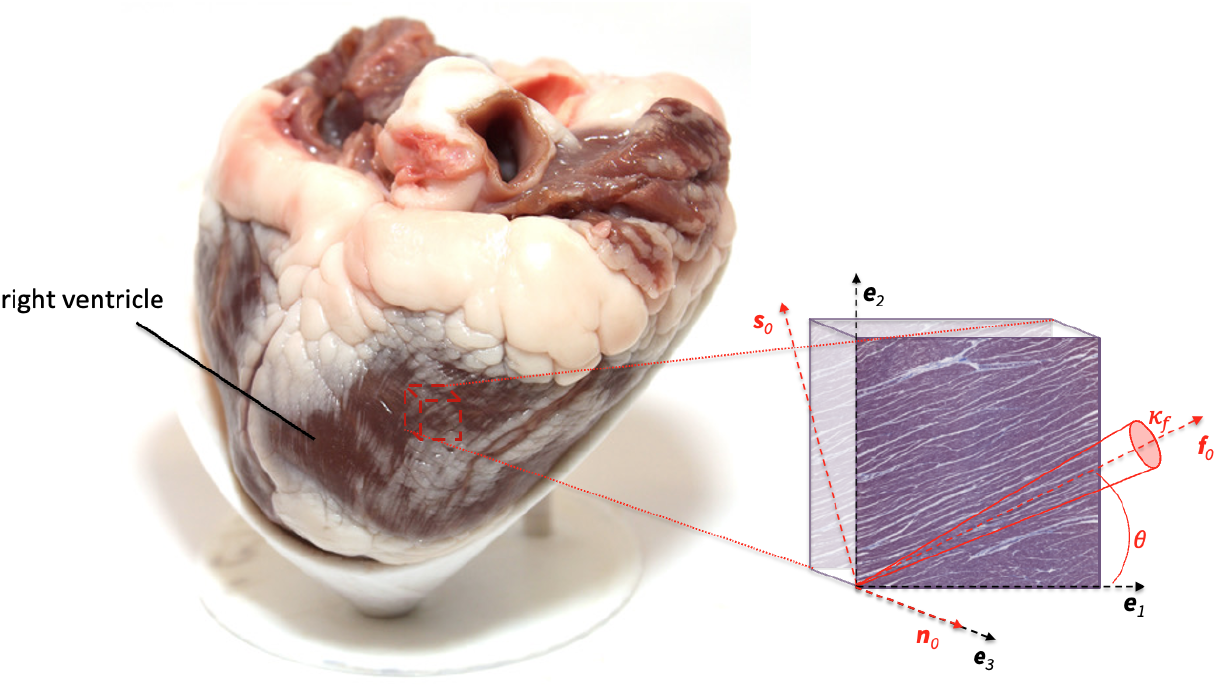
Schematic illustration of the local microstructural coordinate system in the reference configuration in the ovine right ventricle. The fiber (***f***_0_) and sheet (***s***_0_) directions are obtained by a rotation of angle *θ* within the *e*_1_*e*_2_ plane, while the normal direction (***n***_0_) coincides with the *e*_3_ axis. The angle *θ* characterizes the in-plane orientation of the myocardial fibers relative to the global coordinate system. *κ*_*f*_ defines the fiber dispersion in the ***f***_0_ direction.

To account for the orthotropic behavior, we define eight scalar invariants

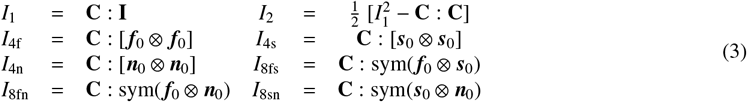

We further account for dispersion in the fiber, sheet, and normal directions (Martonová et al., 2025). For this, we adopt the generalized structure tensor framework (Gasser et al., 2006) and introduce a dispersion parameter, 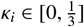 as well as the general structure tensor

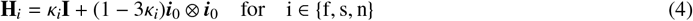

The anisotropic fourth invariants then become

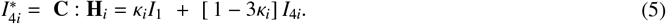

We note that *κ*_*i*_ = 0 corresponds to perfect fiber alignment and 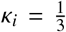 corresponds to the material isotropy. For more details, we refer to the original study (Gasser et al., 2006). To ensure that the fiber and cross-fiber directions can only carry load under tensile deformation while maintaining numerical stability (Holzapfel and Ogden, 2009; Martonová et al., 2024), we employ a smooth activation function instead of the standard max(1, *I*_4*i*_), *i* ∈ {f, s, n} operator and obtain smoothed 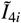. For details, we refer to Appendix A.

Overall, we now consider two distinct types of the microstructural parameters, the mean fiber angle *θ* describing the fiber position in the global coordinate system {***e***_1_, ***e***_2_, ***e***_3_} and three dispersion parameters *κ*_*f*_, *κ*_*s*_, *κ*_*n*_ characterizing the fiber, sheet and fiber-normal dispersions, see Fig. 1. The constitutive response can then be expressed in terms of a strain energy density function

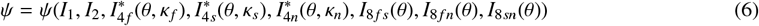

from which we derive the first Piola-Kirchhoff stress as

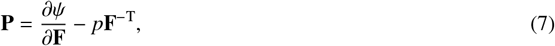

where *p* denotes the hydrostatic pressure, which enforces the assumed incompressibility. This formulation preserves the framework introduced in (Martonová et al., 2025) while extending it to incorporate an in-plane rotation of the mean fiber direction via the parameter *θ*. This extension enables the use of experimental data in which the testing axis does not coincide with the principal fiber and cross-fiber directions, see Section 4.

## 3. Constitutive model discovery

Instead of *a priori* postulating a specific strain energy density function *ψ* as in previous studies of right ventricular myocardium (Kakaletsis et al., 2021; Valdez-Jasso et al., 2012), we identify an appropriate constitutive model using constitutive artificial neural networks (CANNs) (Linka et al., 2021; Linka and Kuhl, 2023; Martonová et al., 2024, 2025).

We formulate the network as a multi-input multi-output (MIMO) framework. The network simultaneously trains on experimental data from multiple testing protocols (e.g. uniaxial tension/compression and triaxial simple shear), as defined in Section 4. Accordingly, it takes multiple strain measures as input and returns the corresponding stress components. However, the network does not map stretches/strains directly to stresses; instead, it approximates the strain energy density function *ψ*, see Figure Fig. 2. From this scalar-valued function, we compute the individual stress components according to Eq. (7) and use them in the optimization problem defined in Eq. (9). For each deformation mode, we construct the architecture to contain the following layers: input layer containing the deformation gradient, invariant layer, dispersion layer, a layer applying polynomial transformations (first and second order), followed by a layer applying identity and exponential functions. Finally, these functions are passed to the shared strain energy layer from which we derive the stress components.

**Figure 2:**
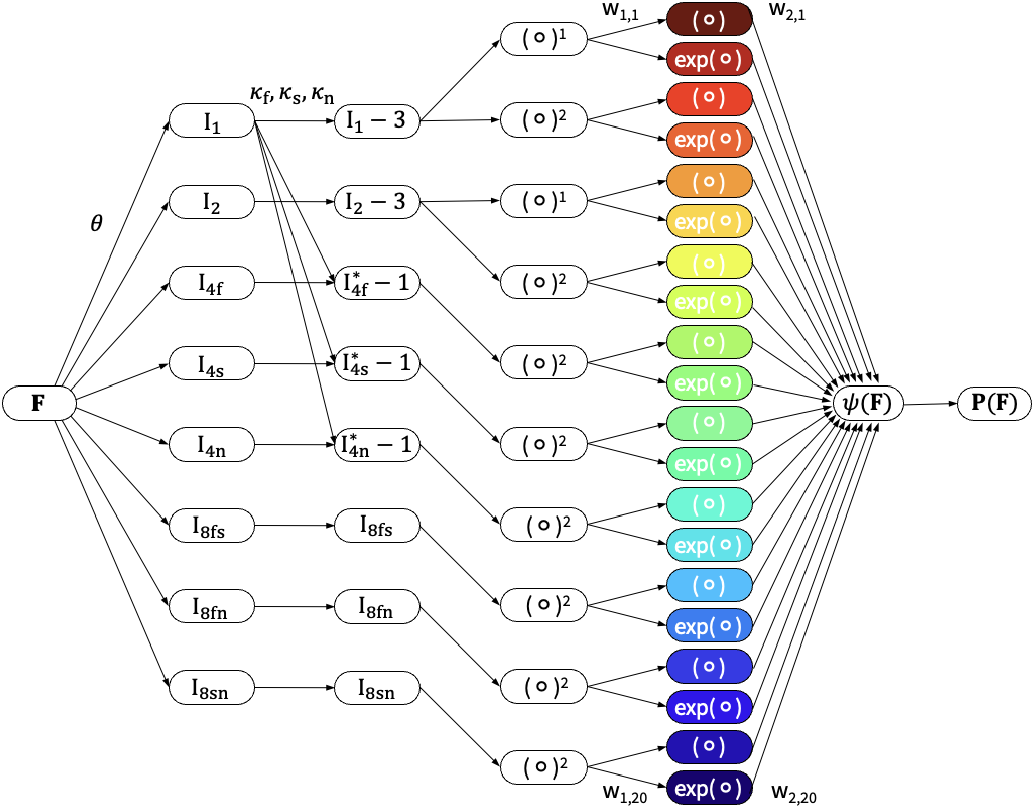
Conceptual architecture of the constitutive artificial neural network. The figure shows the pathway followed by one test protocol. Each test protocol will calculate the deformation gradient **F**, its invariants and stress components based on the continuum model discussed in Section 2. The parameters *θ, κ*_*f*_, *κ*_*s*_ and *κ*_*n*_ are shared among all the test protocols. All protocols pass their invariants through the same instance of strain energy model, hence all contribute simultaneously in learning the set of the weights.

The invariant layer computes eight scalar invariants given in Eq. (3) for each deformation mode specified in Section 4. This defines the multi-input structure of the network. The mean fiber angle *θ* is an optionally trainable parameter.

The dispersion layer incorporates the structural parameters *κ*_*i*_, *i* ∈ {f, s, n}, and computes the dispersed invariants 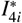 according to Eq. (5). We apply reference shifts by one and three to enforce *ψ*(**F** = **I**) = 0 and share the structural parameters *θ* and *κ*_*i*_ across all testing protocols. This enforces a consistent microstructural representation independent of the applied loading.

The shared strain energy layer is identical for all testing protocols such that the network learns a single constitutive law. To satisfy the requirements of polyconvexity and to guarantee thermodynamic consistency, we constrain all network weights to be non-negative. In contrast to previous works (Martonová et al., 2024, 2025), where the general strain energy function includes both linear and exponential terms for all invariants and the linear contributions of the anisotropic invariants are autonomously driven to zero during training, we omit linear terms associated with anisotropic invariants *a priori* to ensure a stress-free reference configuration (Urrea–Quintero et al., 2026). We retain linear contributions of the isotropic invariants *I*_1_ and *I*_2_. This choice allows us to recover classical models such as the neo-Hookean model, involving the term (*I*_1_ − 3) (Treloar, 1944), the Blatz–Ko model, involving the term (*I*_2_ − 3) (Blatz and Ko, 1962), and the Mooney–Rivlin model (Rivlin, 1947), which combines both terms. These contributions represent established isotropic ground-substance behavior in soft tissue. For anisotropic invariants, Holzapfel-Ogden-type models (Holzapfel and Ogden, 2009; Guan et al., 2019; Martonová et al., 2024) employ quadratic or exponential-quadratic terms to capture the characteristic stiffening of collagen fibers.

From a computational perspective, this design reduces the number of learnable terms from 32 to 20, which improves sparsity and interpretability. The resulting general form of the strain energy density function *ψ* reads

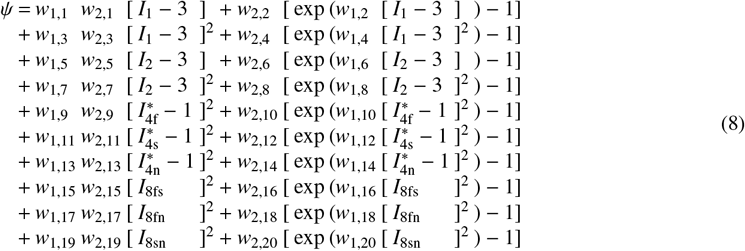

The shared strain energy model outputs the scalar energy density *ψ*. We compute the macroscopic first Piola-Kirchhoff stress **P** through automatic differentiation of *ψ* with respect to the applied stretch or strain, see Eqs. (7), (11) and (12). The resulting stress components correspond to multiple outputs associated with the different testing protocols. Hence, the framework operates as a physics-embedded MIMO model, in which a single set of material and structural parameters simultaneously minimizes the objective function in Eq. (9) for all experimental datasets.

### 3.1. Network training

We train the CANN as a multi-output regression model over nine experimental protocols *p* ∈ 𝒫, six shear and three uniaxial extension tests specified in Section 4, i.e.

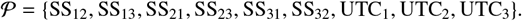

For each protocol *p*, the network predicts the corresponding component of the Piola-Kirchhoff stress *P*_*p,i*_ and we discover the most suitable model and its parameters via minimizing following the objective function

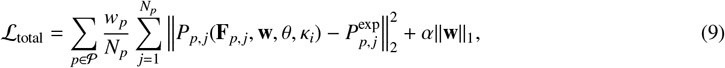

where 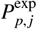 are the experimentally measured stress components, **w** = (*w*_1,1_, …, *w*_2,20_) are the network weights given Eq. (8) and *θ, κ*_*i*_ are optionally trainable microstructural parameters. The normalization factor

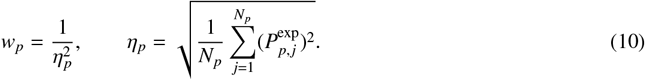

equally scales contributions from protocols with different stress magnitudes (Yong Wang, 2024). To maintain numerical stability for small stress amplitudes, we enforce *η*_*p*_ ≥10^−6^. We also include a *L*_1_-regularization to promote sparsity and encourage the network to identify a parsimonious representation of the strain energy. We set the penalty parameter *α* to 0.01 (Martonová et al., 2026; Vervenne et al., 2025).

We optimize the model using the Adam algorithm (Kingma and Ba, 2017) with a learning rate of 10^−3^, a batch size of 32, and a maximum of 10,000 epochs. We employ a 20% validation split following standard practice(Linka et al., 2021). Because the dataset is ordered by increasing strain, directly applying Keras’ *validation_split* assigns the final portion of the data to the validation set before shuffling^1^. This setup effectively evaluates extrapolation rather than interpolation. To avoid this bias, we shuffle the dataset prior to splitting, which ensures that both subsets follow the same distribution.

We use two callbacks to control the training process and apply an early stopping criterion with a patience of 1000 epochs and a minimum improvement of Δ_min_ = 10^−3^. In addition, *ReduceLROnPlateau*^1^ reduces the learning rate by a factor of 0.5 after 250 epochs without improvement, with a lower bound of 10^−6^. To increase robustness with respect to local minima, we train each sample four times using different initializations of the dispersion parameter *κ*_*f*_ ∈ {0.0, 0.1, 0.2, 0.3}. We then select the model with the highest coefficient of determination (*R*^2^), which improves the stability and reproducibility of the identified material parameters.

## 4. Experimental data

We use experimental data which comprise eleven heart samples from healthy Dorset sheep (Kakaletsis et al., 2021), see Fig. 1. The dataset provides stress-strain data for six simple shear (SS) and stretch–strain data for three uniaxial tension/compression (UTC), all conducted using a triaxial testing system (Sugerman et al., 2021; Jeanpierre et al., 2025). For each deformation mode, we select 64 equidistant data points and use them to train the neural network introduced in Section 3. We assume material incompressibility and consider the following deformation modes characterized by the deformation gradients

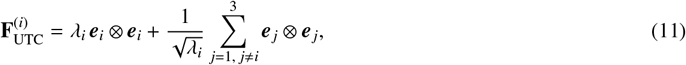

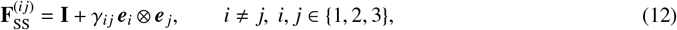

where **I** denotes the identity tensor, *λ*_*i*_ is the stretch in the *i*-direction, and *γ*_*i j*_ is the shear strain. The measurements are given in the global coordinate system {***e***_1_, ***e***_2_, ***e***_3_}.

The data also includes histological information about the mean fiber angle *θ* and the concentration parameter *b*. We convert *b* into the fiber dispersion parameter 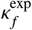 (Gasser et al., 2006), see Tables 1 and 2, column 3, and set the remaining dispersion parameters to *κ*_*s*_ = *κ*_*n*_ = 0.

**Table 1:** Discovered material parameters for right ventricular tissue using dataset 1 (positive and negative stretches and shear strains). Experimental (denoted with superscript exp) and discovered (denoted with superscript *) structural parameters, active invariant sets and mean goodness of fit 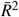 for each individual sample. Results are reported for models with fixed fiber angle *θ*^exp^ and with trainable *θ*. Mean goodness of fit across all samples is 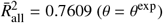 and 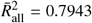 (trainable *θ*).

| Sample | $\theta^{\text{exp}}$ | $\kappa_f^{\text{exp}}$ | $\theta = \theta^{\text{exp}}$ | | | $\theta$ trainable | | | |
| --- | --- | --- | --- | --- | --- | --- | --- | --- | --- |
| | | | $\kappa_f^*$ | $\bar{R}^2$ | active invariants | $\theta^*$ | $\kappa_f^*$ | $\bar{R}^2$ | active invariants |
| 1 | 55.7° | 0.2486 | 0.1431 | 0.8455 | $I_2, I_{4f}, I_{4s}, I_{4n}, I_{8fs}$ | 33.6° | 0.1446 | 0.8710 | $I_1, I_2, I_{4f}, I_{4s}, I_{4n}, I_{8fs}$ |
| 2 | 5.9° | 0.2127 | 0.2352 | 0.8689 | $I_2, I_{4f}, I_{4s}, I_{8fs}, I_{8sn}$ | 4.8° | 0.2327 | 0.8677 | $I_2, I_{4f}, I_{4s}, I_{8fs}, I_{8sn}$ |
| 3 | 41.7° | 0.2108 | 0.3333 | 0.8759 | $I_2, I_{4s}, I_{4n}, I_{8fs}$ | 40.6° | 0.2502 | 0.8765 | $I_2, I_{4s}, I_{4n}, I_{8fs}$ |
| 4 | 6.4° | 0.2274 | 0.1752 | 0.7516 | $I_1, I_2, I_{4f}, I_{4s}, I_{8fs}$ | 0.0° | 0.2036 | 0.8073 | $I_2, I_{4f}, I_{4s}, I_{4n}, I_{8fs}$ |
| 5 | 15.5° | 0.1287 | 0.0627 | 0.6464 | $I_1, I_2, I_{4f}, I_{4s}, I_{4n}, I_{8fn}$ | 7.8° | 0.2280 | 0.6759 | $I_2, I_{4f}, I_{4s}, I_{4n}, I_{8fn}$ |
| 6 | 5.0° | 0.1179 | 0.2087 | 0.7413 | $I_2, I_{4f}, I_{4n}$ | 0.0° | 0.2020 | 0.7494 | $I_2, I_{4f}, I_{4n}$ |
| 7 | 1.7° | 0.0594 | 0.1008 | 0.6953 | $I_2, I_{4f}, I_{4n}, I_{8fs}$ | 0.0° | 0.0636 | 0.7088 | $I_2, I_{4f}, I_{4n}, I_{8fs}$ |
| 8 | 15.6° | 0.2445 | 0.1432 | 0.7733 | $I_2, I_{4f}, I_{4s}, I_{8fs}$ | 0.0° | 0.1711 | 0.7879 | $I_2, I_{4f}, I_{4s}, I_{8fs}$ |
| 9 | 1.3° | 0.1359 | 0.1343 | 0.7488 | $I_1, I_2, I_{4f}, I_{4s}, I_{4n}, I_{8fs}$ | 46.9° | 0.0718 | 0.8051 | $I_2, I_{4f}, I_{8fs}$ |
| 10 | 3.7° | 0.1470 | 0.1889 | 0.8010 | $I_2, I_{4f}, I_{4s}, I_{8fs}$ | 0.0° | 0.2209 | 0.8220 | $I_2, I_{4f}, I_{4s}, I_{8fs}$ |
| 11 | 18.2° | 0.0317 | 0.0000 | 0.6225 | $I_2, I_{4f}, I_{4s}, I_{4n}, I_{8fs}, I_{8fn}$ | 0.0° | 0.1862 | 0.7658 | $I_2, I_{4f}, I_{4n}, I_{8fs}, I_{8fn}$ |

**Table 2:** Discovered material parameters for right ventricular tissue using dataset 2 (positive stretches and shear strains). Experimental (denoted with superscript exp) and discovered (denoted with superscript *) structural parameters, active invariant sets, and mean goodness of fit 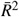 for each sample. Results are reported for models with fixed fiber angle *θ*^exp^ and with trainable *θ*. Mean goodness of fit across all samples is 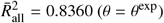 and 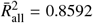 (trainable *θ*).

| Sample | $\theta^{\text{exp}}$ | $\kappa_f^{\text{exp}}$ | $\theta = \theta^{\text{exp}}$ | | | $\theta$ trainable | | | |
| --- | --- | --- | --- | --- | --- | --- | --- | --- | --- |
| | | | $\kappa_f^*$ | $\bar{R}^2$ | active invariants | $\theta^*$ | $\kappa_f^*$ | $\bar{R}^2$ | active invariants |
| 1 | 55.7° | 0.2486 | 0.0000 | 0.7806 | $I_1, I_2, I_{4f}, I_{4s}, I_{4n}, I_{8fs}, I_{8sn}$ | 71.7° | 0.0735 | 0.8817 | $I_1, I_{4f}, I_{4s}, I_{4n}$ |
| 2 | 5.9° | 0.2127 | 0.2634 | 0.8797 | $I_1, I_{4f}, I_{4s}, I_{4n}, I_{8fs}, I_{8sn}$ | 19.1° | 0.0894 | 0.8469 | $I_1, I_2, I_{4f}, I_{4s}, I_{4n}, I_{8sn}$ |
| 3 | 41.7° | 0.2108 | 0.3333 | 0.9195 | $I_1, I_2, I_{4s}, I_{4n}, I_{8fs}$ | 0.0° | 0.0000 | 0.9187 | $I_1, I_{4f}, I_{4s}, I_{4n}$ |
| 4 | 6.4° | 0.2274 | 0.0000 | 0.8611 | $I_1, I_2, I_{4f}, I_{4s}, I_{4n}, I_{8sn}$ | 0.0° | 0.0000 | 0.8922 | $I_1, I_2, I_{4f}, I_{4s}, I_{4n}, I_{8fs}, I_{8fn}$ |
| 5 | 15.5° | 0.1287 | 0.0600 | 0.7113 | $I_1, I_2, I_{4f}, I_{4s}, I_{4n}, I_{8fn}$ | 16.0° | 0.0005 | 0.7319 | $I_2, I_{4f}, I_{4s}, I_{4n}, I_{8fn}$ |
| 6 | 5.0° | 0.1179 | 0.1250 | 0.7935 | $I_1, I_{4f}, I_{4s}, I_{4n}, I_{8fs}$ | 0.0° | 0.0378 | 0.7913 | $I_2, I_{4f}, I_{4s}, I_{4n}, I_{8fs}, I_{8sn}$ |
| 7 | 1.7° | 0.0594 | 0.0000 | 0.8274 | $I_1, I_{4f}, I_{4s}, I_{4n}, I_{8fn}$ | 0.0° | 0.0000 | 0.8293 | $I_1, I_{4f}, I_{4s}, I_{4n}, I_{8fn}$ |
| 8 | 15.6° | 0.2445 | 0.0000 | 0.9328 | $I_1, I_{4f}, I_{4s}, I_{8fs}$ | 15.3° | 0.0000 | 0.9317 | $I_1, I_{4f}, I_{4s}, I_{8fs}$ |
| 9 | 1.3° | 0.1359 | 0.2222 | 0.8641 | $I_1, I_{4f}, I_{4s}, I_{8fs}$ | 13.8° | 0.1407 | 0.8684 | $I_1, I_{4f}, I_{4s}$ |
| 10 | 3.7° | 0.1470 | 0.0000 | 0.9049 | $I_1, I_{4f}, I_{4s}, I_{4n}, I_{8fs}, I_{8sn}$ | 0.0° | 0.0000 | 0.9161 | $I_1, I_{4f}, I_{4s}, I_{4n}, I_{8fs}, I_{8sn}$ |
| 11 | 18.2° | 0.0317 | 0.0000 | 0.7214 | $I_1, I_{4f}, I_{4s}, I_{4n}, I_{8fs}, I_{8fn}$ | 0.0° | 0.0000 | 0.8439 | $I_1, I_2, I_{4f}, I_{4s}, I_{4n}, I_{8fs}, I_{8fn}$ |

## 5. Results and Discussion

We evaluate the performance of the constitutive neural network under four learning scenarios designed to assess both predictive accuracy and the identifiability of structural parameters. Tables 1 and 2 as well as B.3 and B.4 summarize the results for dataset 1 and dataset 2, respectively. Dataset 1 contains uniaxial tension and compression together with simple shear over the full strain range, including both positive and negative values. Dataset 2 is restricted to positive stretches and shear strains only. For each dataset, we compare two configurations in which the fiber angle *θ* remains fixed to its histological measurement *θ*^exp^ or acts as a trainable parameter.

### 5.1. Discovered models and active invariants

The discovered models consistently rely on a compact but expressive set of invariants. The isotropic invariant *I*_2_ and the fiber-related invariant *I*_4f_ appear in most samples, see Tables 2 and B.3 for the sample-specific strain energy functions. It confirms that both the matrix and the fiber contributions are essential to describe the mechanical response. In addition, the model frequently activates anisotropic invariants such as *I*_4s_ and *I*_4n_, as well as mixed invariants like *I*_8fs_, *I*_8fn_, and *I*_8sn_. These mixed terms capture interactions between fiber families and shear deformation and play a particularly important role in samples with pronounced anisotropic coupling.

A notable observation is that several samples, in particular in dataset 2 (see Table 2), recover invariant combinations that resemble the classical Holzapfel-Ogden model (Holzapfel and Ogden, 2009). In particular, the discovered models for sample 8 (both configurations), as well as for sample 9 (fixed *θ*^exp^) involve *I*_1_ together with the fourth invariants *I*_4f_ and *I*_4s_, and the coupling invariant *I*_8fs_, which shows that the network identifies physically meaningful and well-established constitutive models.

### 5.2. Training on dataset 1 versus dataset 2

Training on dataset 1 with fixed fiber angle *θ*^exp^ yields an average coefficient of determination of 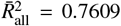 across all samples and loading modes. Table 1 summarizes the results for the best-performing run of each sample. The identified invariant sets frequently include coupling terms such as *I*_8fs_ and *I*_8fn_, which reflects the richer mechanical information contained in the full strain range. The predicted dispersion parameters *κ*_*f*_ show good agreement with the experimentally measured values, as illustrated in Fig. 3 (left), where the predictions cluster closely around the line of perfect agreement.

**Figure 3:**
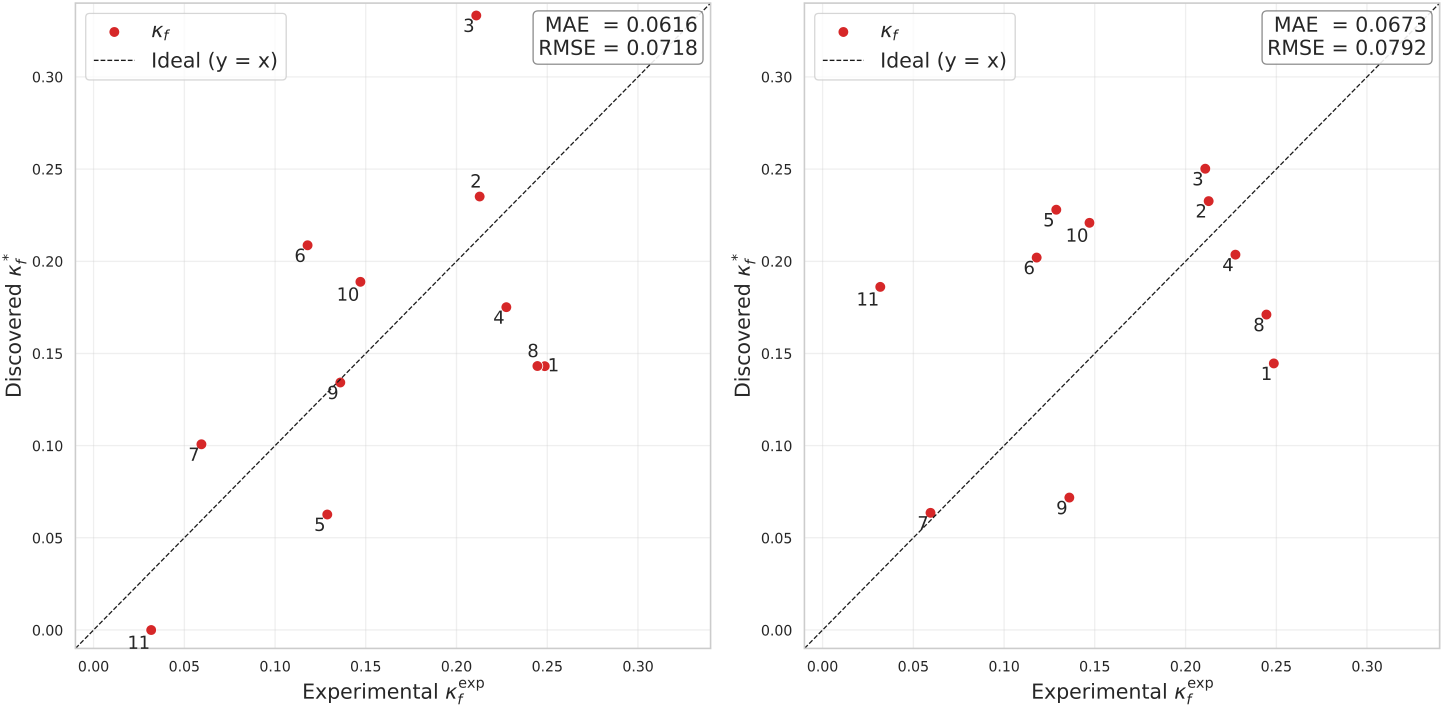
Comparison of experimental 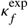 and discovered 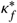values using the CANN model for dataset 1 for each sample. Each scatter plot shows the discovered versus experimental *κ*_*f*_, with the dashed line indicating ideal agreement (*y* = *x*). Left: model trained with a fixed fiber angle *θ* = *θ*^exp^. Right: model trained with *θ* as a trainable parameter. Mean absolute error (MAE) and root mean square error (RMSE).

As a representative example, Fig. 4 shows the predicted and experimental stress response for sample 9. The model accurately reproduces the mechanical response across simple shear and uniaxial loading modes and recovers the dispersion parameter *κ*_*f*_ with high accuracy, with a deviation of approximately 1% from the experimental value (see Table 1 left).

**Figure 4:**
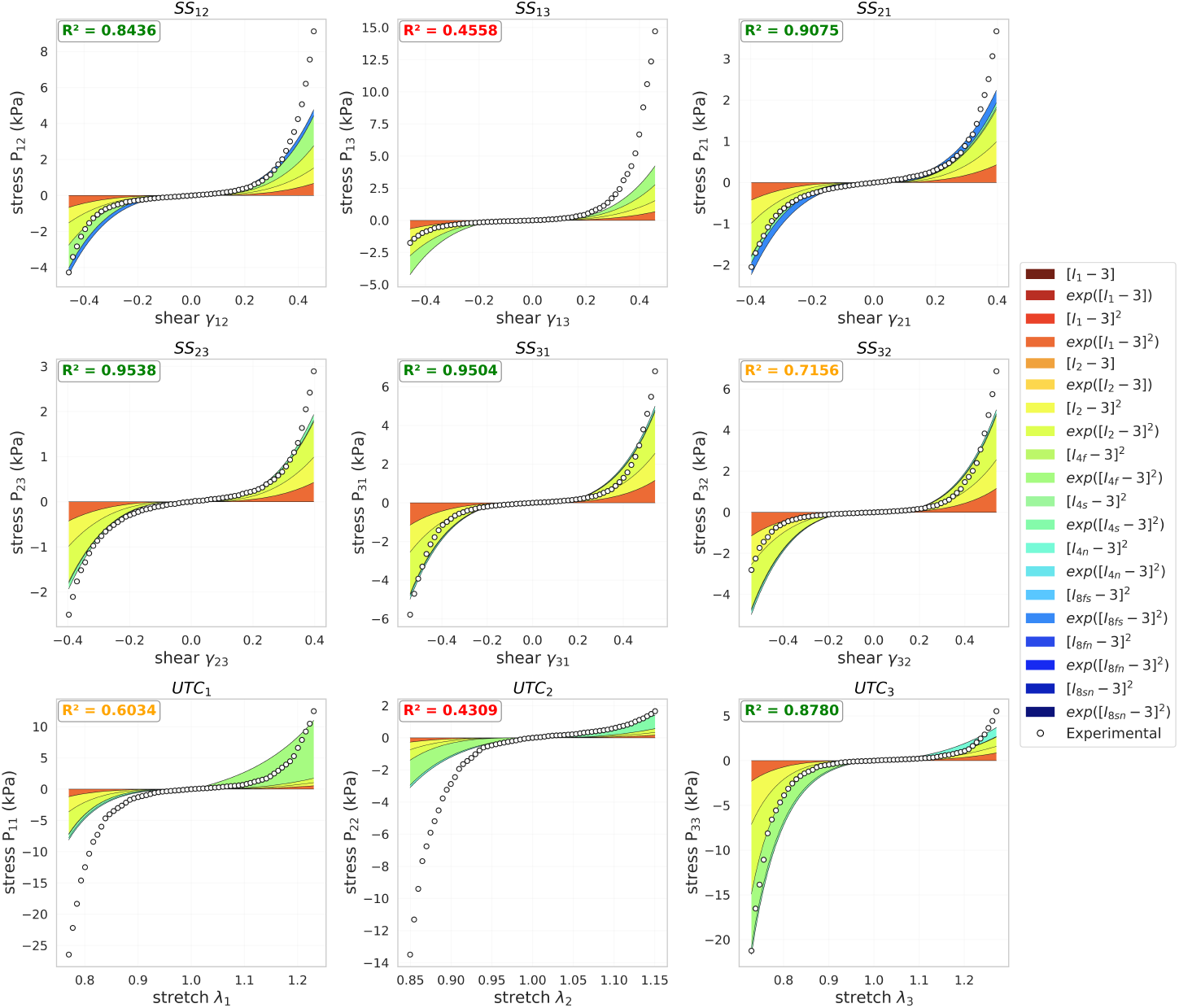
Model discovery for Sample 9 trained on positive and negative strain values. First Piola-Kirchhoff stress components as functions of shear strain during simple shear tests (SS), first and second row, and stretches during uniaxial tension and compression tests (UTC), third row, for orthotropic, perfectly incompressible CANN from Fig. 2. Dots represent experimental data; color-coded areas highlight the 20 term contributions in the discovered stress function. Discovered 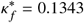, mean 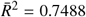for all testing modes.

Allowing the fiber angle *θ* to vary during training increases the average coefficient of determination to 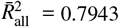. The model uses the additional flexibility to compensate for discrepancies between the histological fiber orientation and the effective mechanical response. However, this improvement in goodness of fit comes at the expense of physical interpretability. As shown in Fig. 3 right, the predictions of the dispersion parameter *κ*_*f*_ exhibit larger deviations from the line of perfect agreement, indicating reduced accuracy in *κ*_*f*_ identification. This behavior suggests that the model compensates for structural mismatches by adjusting the fiber orientation instead of consistently identifying the underlying microstructural parameters. In some cases, the model also alters the balance between invariant contributions, which reduces the uniqueness of the identified parameters.

Restricting the data to positive strains (dataset 2) leads to a higher predictive accuracy. This restriction reflects a common experimental and modeling practice, since collagen fibers primarily carry load in tension while exhibiting negligible stiffness in compression. With fixed fiber angle, the average coefficient of determination increases to 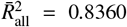. Table 2 summarizes these results and Fig. 6 exemplarily shows the stress–strain (SS) and stress–stretch (UTC) curves for sample 9. The model achieves improved agreement with the experimental data across all loading modes. The structure of the discovered models changes in this setting. The invariant sets become simpler and often resemble classical formulations based on *I*_1_ and fiber invariants. The model uses fewer coupling terms compared to dataset 1. This result indicates that the reduced dataset does not provide sufficient information to activate more complex interaction mechanisms.

Despite the improved predictive accuracy, the identification of the dispersion parameter *κ*_*f*_ deteriorates. Several samples show *κ*_*f*_ = 0 or strong deviations from the experimental values, as reported in Table 2 and Fig. 5. This result indicates that fiber dispersion cannot be uniquely identified from tensile data alone.

**Figure 5:**
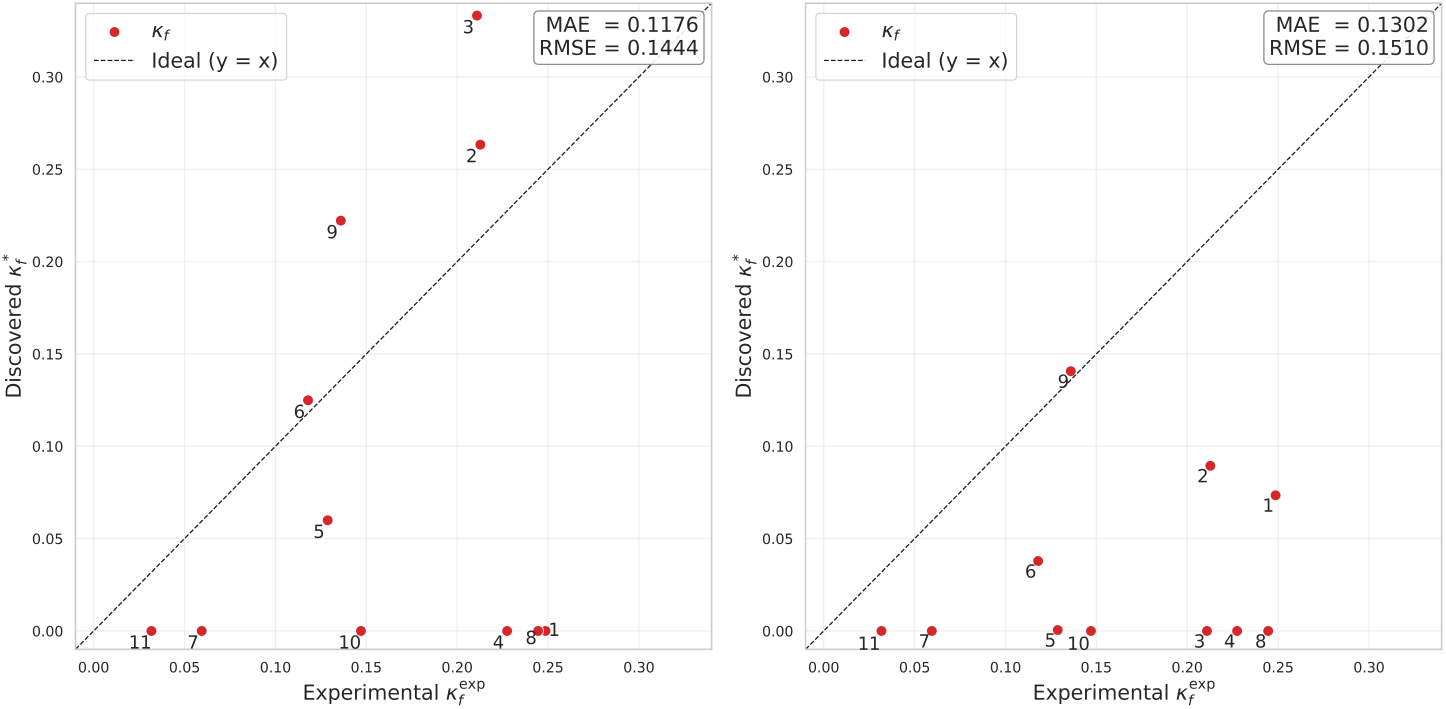
Comparison of experimental 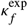 and discovered 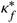 values using the CANN model for dataset 2 for each sample. Each scatter plot shows the discovered versus experimental *κ*_*f*_, with the dashed line indicating ideal agreement (*y* = *x*). Left: model trained with a fixed fiber angle *θ* = *θ*^exp^. Right: model trained with *θ* as a trainable parameter. Mean absolute error (MAE) and root mean square error (RMSE).

**Figure 6:**
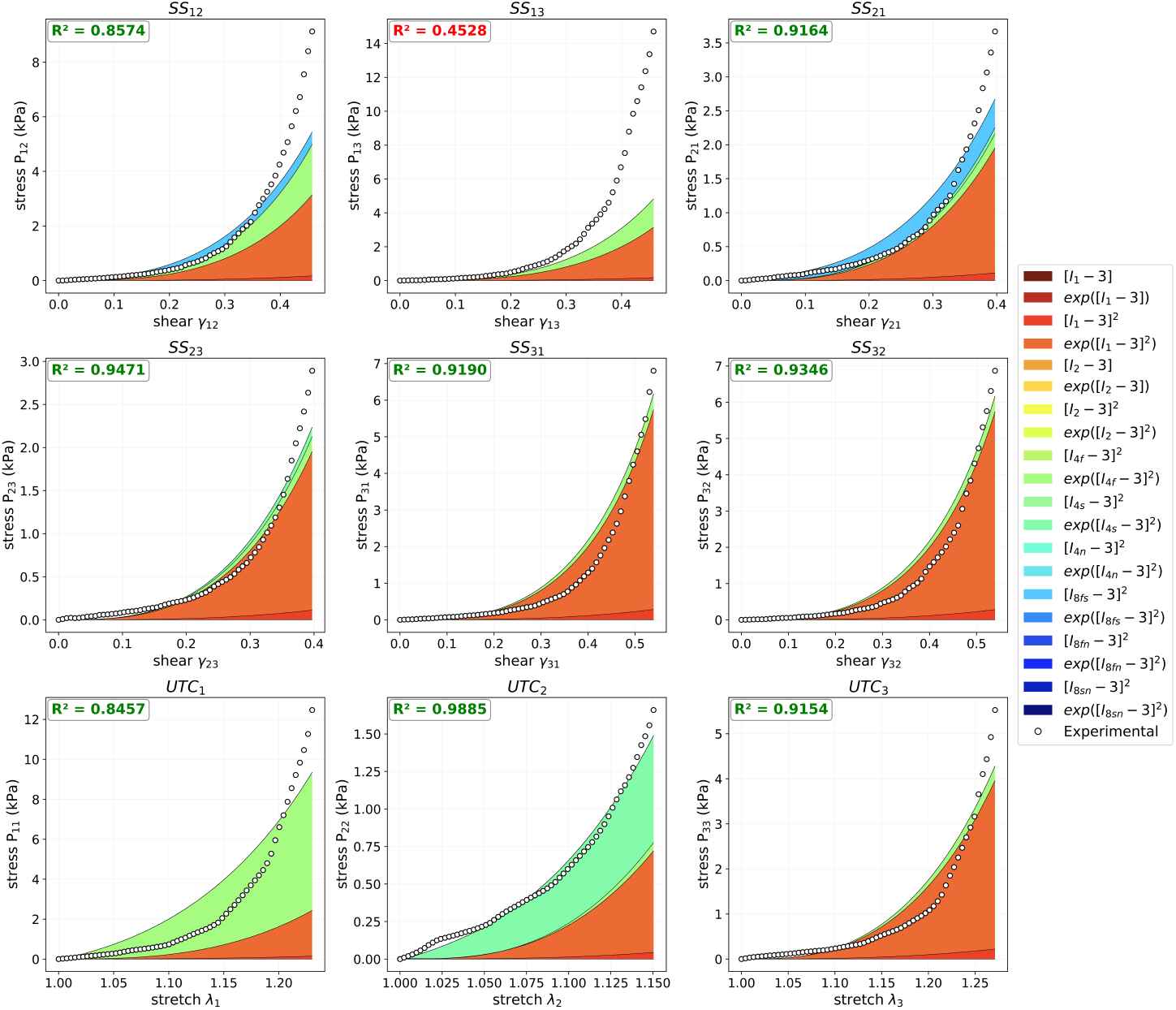
Model discovery for sample 9 trained on positive strain values. First Piola-Kirchhoff stress components as functions of shear strain during simple shear tests (SS), first and second row, and stretches during uniaxial tension and compression tests (UTC), third row, for orthotropic, perfectly incompressible CANN from Fig. 2. Dots represent experimental data; color-coded areas highlight the 20 term contributions in the discovered stress function. Discovered 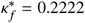, mean 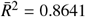 for all loading modes.

### 5.3. Trade-off between predictive accuracy and parameter identifiability

The comparison between dataset 1 and dataset 2 reveals a clear trade-off between predictive accuracy and parameter identifiability. Dataset 2 produces higher *R*^2^ values; however, this improvement does not reflect a better physical description of the material. Instead, it highlights the fundamental difference between phenomenological curve-fitting and physically meaningful parameter identification.

This behavior originates from the tension-compression asymmetry of collagen fibers. Fibers carry load in tension but buckle under compression, and this mechanism is explicitly enforced in the CANN architecture through a tension-only activation of the fiber invariants, see Appendix A. In dataset 1, the inclusion of compressive strain regimes suppresses the fiber contribution and isolates the response of the isotropic ground matrix. This separation enables the model to identify the matrix behavior independently and attribute the remaining stiffening in tension to the fiber network, which leads to accurate recovery of the dispersion parameter *κ*_*f*_.

In contrast, dataset 2 contains only tensile loading, where both matrix and fiber contributions remain active and strongly coupled. This coupling renders the parameter identification problem ill-posed, since multiple combinations of parameters can produce similar stress responses. As a result, the network achieves higher *R*^2^ values by compensating for incorrect dispersion parameters through adjustments of other invariant contributions. Consequently, the restriction to positive strains simplifies the optimization problem but reduces the ability to uniquely identify the underlying microstructural parameters.

Our results demonstrate that high predictive accuracy alone does not guarantee physically meaningful parameter identification. Reliable identification requires sufficiently rich kinematic data that enables separation of the underlying mechanical mechanisms.

Some limitations warrant discussion. First, we assume full tissue incompressibility and therefore may not capture the slight compressibility of myocardial tissue (Bonnemains et al., 2019; Avazmohammadi et al., 2020). Second, we represent fiber dispersion only through the fourth invariants and thus may miss more complex anisotropic and coupling effects (Melnik et al., 2018). Third, we train the model on a limited dataset and restrict it to passive behavior, which limits generalizability and calls for extensions to larger datasets and active mechanics.

## 6. Conclusion

We present a physics-embedded constitutive neural network for automated discovery of anisotropic material models in right ventricular myocardium. The results show that inclusion of compressive deformation modes is essential for reliable identification of microstructural parameters such as fiber dispersion, whereas training on tensile data alone increases predictive accuracy but reduces parameter identifiability. The proposed framework enables model discovery directly from individual experimental right ventricular data and identifies both classical invariant-based structures and microstructural parameters such as fiber orientation and dispersion.

## Code and data availability

Upon final publication, our source code, data, and examples will be available online.

## Acknowledgments

The authors acknowledge support from the European Research Council (ERC) Grant 101141626 DISCOVER funded by the European Union to EK, from FAU Emerging Talents Initiative (ETI) to DM. Views and opinions expressed are, however, those of the authors only and do not necessarily reflect those of the European Union or the European Research Council Executive Agency. Neither the European Union nor the granting authority can be held responsible for them.

## Appendix A. Exclusion of compressed fibers

The smoothed active invariant 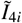, *i* ∈ {f, s, n} is defined using a scaled *softplus* formulation:

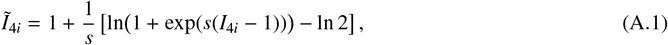

where *s* is a sharpness hyperparameter (e.g., *s* = 100). The first derivative of this activation, which directly scales the fiber contribution to the macroscopic stress, yields a smooth sigmoid function:

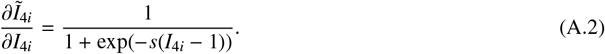

Although modern automatic differentiation frameworks can compute subgradients for non-smooth functions such as the maximum function, we deliberately avoid its use due to its lack of 𝒞^2^-continuity, which is essential for robust continuum mechanics modeling. The max operator introduces a discontinuous Heaviside step in its first derivative, leading to an abrupt and non-physical onset of stress that does not reflect the progressive uncrimping of collagen fibers. More critically, its second derivative corresponds to a Dirac delta distribution, resulting in an ill-defined material tangent stiffness tensor. This often leads to convergence issues in Newton-Raphson iterations within macroscopic finite element method (FEM) solvers. In contrast, our proposed smooth approximation guaranties physically realistic stress evolution and a well-defined, continuous stiffness matrix, enabling stable and reliable integration into FEM simulations.

Crucially, this activation is applied directly to the raw invariant *I*_4*i*_ prior to dispersion mixing to ensure physical consistency and numerical stability. Although the standard Gasser-Ogden-Holazpfel (GOH) model (Gasser et al., 2006) traditionally applies a macroscopic switch to the dispersed invariant, this has been shown to produce non-physical results - specifically, the total loss of fiber contribution when the mean direction is compressed, even if off-axis fibers remain in tension (Latorre Marcos, 2016). By adopting a “pre-integrated” logic where individual fiber buckling is handled via the smooth fiber activation before dispersion, the model allows the *κ*_*i*_*I*_1_ term to maintain the load-bearing capacity of dispersed fibers. This sequence aligns with the microstructural reality of collagen behavior and ensures *C*^1^ continuity throughout the tension-compression transition.

## Appendix B. Discovered strain energy functions for right ventricular samples

**Table B.3:** Discovered strain energy function *ψ* and parameters for right ventricular tissue using dataset 1 (positive and negative stretches and shear strains and experimentally measured mean fiber angle *θ*^exp^.

| sample | discovered strain energy function $\psi$ |
| --- | --- |
| 1 | $1.84[I_2 - 3]^2 + 2.12[\exp(2.49[I_2 - 3]^2) - 1] + 3.86[I_{4f}^* - 1]^2 + 0.24[\exp(3.33[I_{4s}^* - 1]^2) - 1] + 0.64[I_{4n}^* - 1]^2 + 1.73[\exp(2.02[I_{8fs}]^2) - 1]$ |
| 2 | $1.79[\exp(1.77[I_2 - 3]^2) - 1] + 17.29[I_{4f}^* - 1]^2 + 4.20[\exp(4.08[I_{4f}^* - 1]^2) - 1] + 2.39[I_{4s}^* - 1]^2 + 0.79[\exp(0.73[I_{4s}^* - 1]^2) - 1] + 0.58[\exp(1.10[I_{8fs}]^2) - 1] + 0.54[\exp(0.75[I_{8sn}]^2) - 1]$ |
| 3 | $1.32[I_2 - 3]^2 + 2.32[\exp(2.93[I_2 - 3]^2) - 1] + 0.75[\exp(0.87[I_{4s}^* - 1]^2) - 1] + 0.78[\exp(0.98[I_{4n}^* - 1]^2) - 1] + 2.63[I_{8fs}]^2$ |
| 4 | $0.66[\exp(1.14[I_1 - 3]^2) - 1] + 0.38[I_2 - 3] + 1.26[\exp(1.83[I_2 - 3]^2) - 1] + 11.72[I_{4f}^* - 1]^2 + 3.59[\exp(3.39[I_{4f}^* - 1]^2) - 1] + 1.59[I_{4s}^* - 1]^2 + 0.52[\exp(0.64[I_{8fs}]^2) - 1]$ |
| 5 | $0.01[I_1 - 3]^2 + 0.05[\exp(0.11[I_1 - 3]^2) - 1] + 0.35[\exp(0.23[I_2 - 3]) - 1] + 0.93[\exp(1.14[I_2 - 3]^2) - 1] + 1.61[I_{4f}^* - 1]^2 + 1.55[\exp(1.50[I_{4f}^* - 1]^2) - 1] + 0.14[I_{4s}^* - 1]^2 + 0.55[\exp(0.48[I_{4s}^* - 1]^2) - 1] + 0.88[I_{4n}^* - 1]^2 + 0.84[\exp(0.79[I_{4n}^* - 1]^2) - 1] + 0.27[I_{8fn}]^2 + 0.77[\exp(0.85[I_{8fn}]^2) - 1]$ |
| 6 | $1.84[\exp(2.55[I_2 - 3]^2) - 1] + 9.33[I_{4f}^* - 1]^2 + 0.69[I_{4n}^* - 1]^2$ |
| 7 | $0.98[I_2 - 3]^2 + 1.38[\exp(1.86[I_2 - 3]^2) - 1] + 3.01[I_{4f}^* - 1]^2 + 1.75[\exp(1.16[I_{4f}^* - 1]^2) - 1] + 0.66[\exp(1.00[I_{4n}^* - 1]^2) - 1] + 0.11[I_{8fs}]^2$ |
| 8 | $1.99[\exp(2.33[I_2 - 3]^2) - 1] + 2.09[I_{4f}^* - 1]^2 + 2.37[\exp(2.20[I_{4f}^* - 1]^2) - 1] + 0.72[I_{4s}^* - 1]^2 + 0.85[\exp(0.83[I_{4s}^* - 1]^2) - 1] + 0.55[\exp(1.19[I_{8fs}]^2) - 1]$ |
| 9 | $1.24[\exp(1.33[I_1 - 3]^2) - 1] + 2.23[I_2 - 3]^2 + 1.92[\exp(1.56[I_2 - 3]^2) - 1] + 2.98[\exp(2.35[I_{4f}^* - 1]^2) - 1] + 0.73[\exp(0.72[I_{4s}^* - 1]^2) - 1] + 0.33[I_{4n}^* - 1]^2 + 0.47[\exp(0.71[I_{8fs}]^2) - 1]$ |
| 10 | $2.19[I_2 - 3]^2 + 2.05[\exp(1.65[I_2 - 3]^2) - 1] + 7.04[I_{4f}^* - 1]^2 + 3.44[\exp(3.15[I_{4f}^* - 1]^2) - 1] + 0.97[I_{4s}^* - 1]^2 + 0.46[I_{8fs}]^2$ |
| 11 | $1.31[\exp(1.37[I_2 - 3]^2) - 1] + 1.89[I_{4f}^* - 1]^2 + 0.55[\exp(0.62[I_{4s}^* - 1]^2) - 1] + 1.10[I_{4n}^* - 1]^2 + 0.55[\exp(0.82[I_{8fs}]^2) - 1] + 0.65[\exp(1.34[I_{8fn}]^2) - 1]$ |

**Table B.4:** Discovered strain energy function *ψ* and parameters for right ventricular tissue using dataset 2 (positive stretches and shear strains) and experimentally measured mean fiber angle *θ*^exp^.

| sample | discovered strain energy function $\psi$ |
| --- | --- |
| 1 | $0.26[\exp(0.23[I_1 - 3]) - 1] + 2.18[I_1 - 3]^2 + 1.80[\exp(1.60[I_1 - 3]^2) - 1] + 0.92[\exp(0.87[I_2 - 3]^2) - 1] + 0.99[\exp(0.84[I_{4f}^* - 1]^2) - 1] + 1.30[\exp(1.39[I_{4s}^* - 1]^2) - 1] + 0.47[\exp(1.14[I_{4n}^* - 1]^2) - 1] + 0.86[I_{8fs}]^2 + 1.04[\exp(1.02[I_{8fs}]^2) - 1] + 0.02[I_{8sn}]^2$ |
| 2 | $0.67[I_1 - 3]^2 + 18.75[I_{4f}^* - 1]^2 + 4.95[\exp(5.25[I_{4f}^* - 1]^2) - 1] + 1.06[\exp(1.83[I_{4s}^* - 1]^2) - 1] + 0.24[I_{4n}^* - 1]^2 + 0.37[\exp(3.06[I_{8fs}]^2) - 1] + 0.82[I_{8sn}]^2$ |
| 3 | $2.09[\exp(3.22[I_1 - 3]^2) - 1] + 0.25[\exp(0.79[I_2 - 3]^2) - 1] + 1.67[\exp(1.81[I_{4s}^* - 1]^2) - 1] + 0.65[\exp(1.04[I_{4n}^* - 1]^2) - 1] + 0.73[\exp(1.40[I_{8fs}]^2) - 1]$ |
| 4 | $0.74[\exp(1.65[I_1 - 3]^2) - 1] + 1.13[\exp(2.18[I_2 - 3]^2) - 1] + 1.82[\exp(2.38[I_{4f}^* - 1]^2) - 1] + 1.34[\exp(1.95[I_{4s}^* - 1]^2) - 1] + 0.66[\exp(0.62[I_{4n}^* - 1]^2) - 1] + 0.25[\exp(1.35[I_{8sn}]^2) - 1]$ |
| 5 | $0.93[\exp(1.01[I_1 - 3]^2) - 1] + 0.09[\exp(0.18[I_2 - 3]^2) - 1] + 0.87[I_{4f}^* - 1]^2 + 1.32[\exp(1.60[I_{4f}^* - 1]^2) - 1] + 0.41[\exp(1.39[I_{4s}^* - 1]^2) - 1] + 0.49[\exp(0.78[I_{4n}^* - 1]^2) - 1] + 0.60[\exp(1.06[I_{8fn}]^2) - 1]$ |
| 6 | $1.07[I_1 - 3]^2 + 1.35[\exp(1.59[I_1 - 3]^2) - 1] + 2.86[I_{4f}^* - 1]^2 + 0.39[I_{4s}^* - 1]^2 + 0.09[\exp(2.34[I_{4n}^* - 1]^2) - 1] + 0.07[I_{8fs}]^2$ |
| 7 | $1.26[\exp(1.94[I_1 - 3]^2) - 1] + 0.02[\exp(9.70[I_{4f}^* - 1]^2) - 1] + 0.26[\exp(1.86[I_{4s}^* - 1]^2) - 1] + 0.35[I_{4n}^* - 1]^2 + 0.23[\exp(0.81[I_{8fn}]^2) - 1]$ |
| 8 | $1.67[\exp(2.53[I_1 - 3]^2) - 1] + 0.34[\exp(4.53[I_{4f}^* - 1]^2) - 1] + 0.54[\exp(0.65[I_{4s}^* - 1]^2) - 1] + 0.30[\exp(2.93[I_{8fs}]^2) - 1]$ |
| 9 | $0.45[I_1 - 3]^2 + 2.41[\exp(2.84[I_1 - 3]^2) - 1] + 3.40[\exp(4.04[I_{4f}^* - 1]^2) - 1] + 0.63[\exp(0.72[I_{4s}^* - 1]^2) - 1] + 0.52[I_{8fs}]^2$ |
| 10 | $1.83[\exp(2.70[I_1 - 3]^2) - 1] + 2.75[I_{4f}^* - 1]^2 + 0.57[\exp(0.92[I_{4s}^* - 1]^2) - 1] + 0.38[\exp(0.52[I_{4n}^* - 1]^2) - 1] + 0.24[\exp(1.73[I_{8fs}]^2) - 1] + 0.12[I_{8sn}]^2$ |
| 11 | $1.24[\exp(1.62[I_1 - 3]^2) - 1] + 0.54[I_{4f}^* - 1]^2 + 0.53[\exp(0.51[I_{4s}^* - 1]^2) - 1] + 0.31[\exp(0.81[I_{4n}^* - 1]^2) - 1] + 0.31[\exp(1.22[I_{8fs}]^2) - 1] + 0.70[\exp(0.84[I_{8fn}]^2) - 1]$ |

## Footnotes

1 Keras `model.fit` documentation: https://keras.io/api/models/model_training_apis/

